# Cerebellum violates Marr-Albus predictions to train synapses on long-term anticipatory goals

**DOI:** 10.64898/2026.04.21.719977

**Authors:** Christian Hansel, Ting-Feng Lin

## Abstract

The cerebellar role in motor adaptation rests – according to the foundational theories of Marr and Albus – on the detection of (near)coincident activity between parallel fiber (PF) and climbing fiber (CF) inputs onto Purkinje cells. Supported by numerous *in vitro* studies, these theories predict synaptic adaptation based on temporal precision detected in time windows of 0ms to ∼100ms. These predictions have not been tested in intact animals. Using two-photon imaging from cerebellar Crus I in intact, awake mice, we show here that coincident stimulation of the PF and CF inputs does not initiate plasticity. Rather, long-term depression (LTD) is reliably evoked by ramping activity of the PF pathway that precedes a CF burst by 400ms. These observations demonstrate a cerebellar plasticity that is not about precision in coincidence with CF signaling. Rather, it shows that cerebellar learning centers on the evaluation of anticipatory PF signals by the CF input.

## Introduction

Climbing fibers (CFs) play a crucial role in the ability of the cerebellum to fine-tune movements. The short temporal delay between parallel fiber (PF) and CF-mediated signals onto Purkinje cells (PCs)^1^ motivated a strong emphasis on coincident signaling in Marr-Albus theories of cerebellar plasticity and learning^2,3^. Recordings from decerebrate rabbits confirmed that co-activation of vestibular mossy fibers (ultimately causing PF activation) and the CF input would initiate a long-term depression (LTD) of PF-PC synapses^4^, but the continuous vestibular stimulation at 20Hz and inferior olive stimulation at 4Hz prevented an analysis of critical temporal relationships between the PF and CF pulses. These relationships were subsequently characterized in numerous *in vitro* studies^5-8^, in which recordings from cerebellar slices confirmed that LTD was induced upon PF+CF activation at short intervals characterized by Marr as ‘*at about the same time*’ (up to ∼100ms interval duration)^2^.

A synaptic interaction at a fundamentally different time scale is a CF burst that is preceded by ramping activity at PF synapses. ‘Ramping’ describes an increase in activity that takes place over hundreds of milliseconds to seconds and has been observed in granule cells and PCs, for example in anticipation of cues to trigger learned behaviors (for ramping in different contexts^9-12^). Coincident and prolonged, preceding PF activity patterns, respectively, signify different relationships between PF and CF inputs. The former marks the type of temporal precision that Marr considered necessary to allow the cerebellum to learn motor skills^2^. The latter, in contrast, better corresponds with modern theories of a cerebellar role in predictive coding^13,14^.

Here, we asked whether in the cerebellum of intact, awake mice both coincidence and ramping synaptic activation protocols would be equally efficient to drive LTD of PF responses. We acknowledge that multiple forms of plasticity operate at different sites in the cerebellar circuit^15^, but here intentionally focus on a specific form of synaptic plasticity that results from the interaction between the two excitatory input types received by PCs. CF activation causes complex spike firing in PCs and subsequently widespread calcium transients^16^. These signals reach supralinear levels when the PF is coactivated^17^ and define a calcium threshold rule, in which the threshold for LTD induction is higher than that for LTP^7^. Would the LTD induction rules established in brain slices equally apply in the intact, awake brain? On the other hand, would ramping activity in the PF pathway differentially trigger LTD?

To determine cerebellar plasticity rules *in vivo*, we used two-photon recordings from awake mice. We found that prolonged patterns of PF activity preceding a CF burst efficiently drive LTD at PF to PC synapses. In contrast, LTD was not observed when classic PF+CF stimulation protocols were applied, in which pulses from the two input pathways were delivered coincidently. By further analyzing calcium dynamics during the induction protocols, we observed that a successful LTD outcome did not depend on the peak amplitude of evoked calcium transients, but rather on the prolonged calcium buildup during PF inputs, which might be critical to set the calcium threshold for LTD induction.

## Results

### CF stimulation elicits calcium transients in PF spines on fine dendritic branches

To assess PF synaptic plasticity, we performed two-photon imaging experiments from cerebellar Crus I of awake mice. jGCaMP7b was specifically expressed in PCs by co-injection of AAV vectors encoding cre-recombinase under control of the Purkinje cell protein 2 (Pcp2)/L7 promoter and Cre-dependent jGCaMP7b (Fig. 1a). Electrical stimulation was introduced using glass pipettes filled with artificial cerebrospinal fluid (ACSF). Electrodes were located (1) in the molecular layer at least 100 μm lateral from the recorded PCs to stimulate PFs and (2) in the granule cell layer below the recorded PCs to stimulate the CFs (Fig. 1a). PC dendrites expressing jGCaMP7b were identified in the top view as thin stripes with a sagittal orientation (Fig. 1b). PF stimulation (8 pulses at 100Hz) caused a beam-like pattern of activated PC dendrites, while the same stimulus pattern applied to the CFs caused a sagittal activation pattern oriented perpendicular to the PF beam (Fig. 1c). In individual PC dendrites, stimulus-locked responses were seen with both PF and CF stimulation (Fig. 1d).

**Fig. 1.**
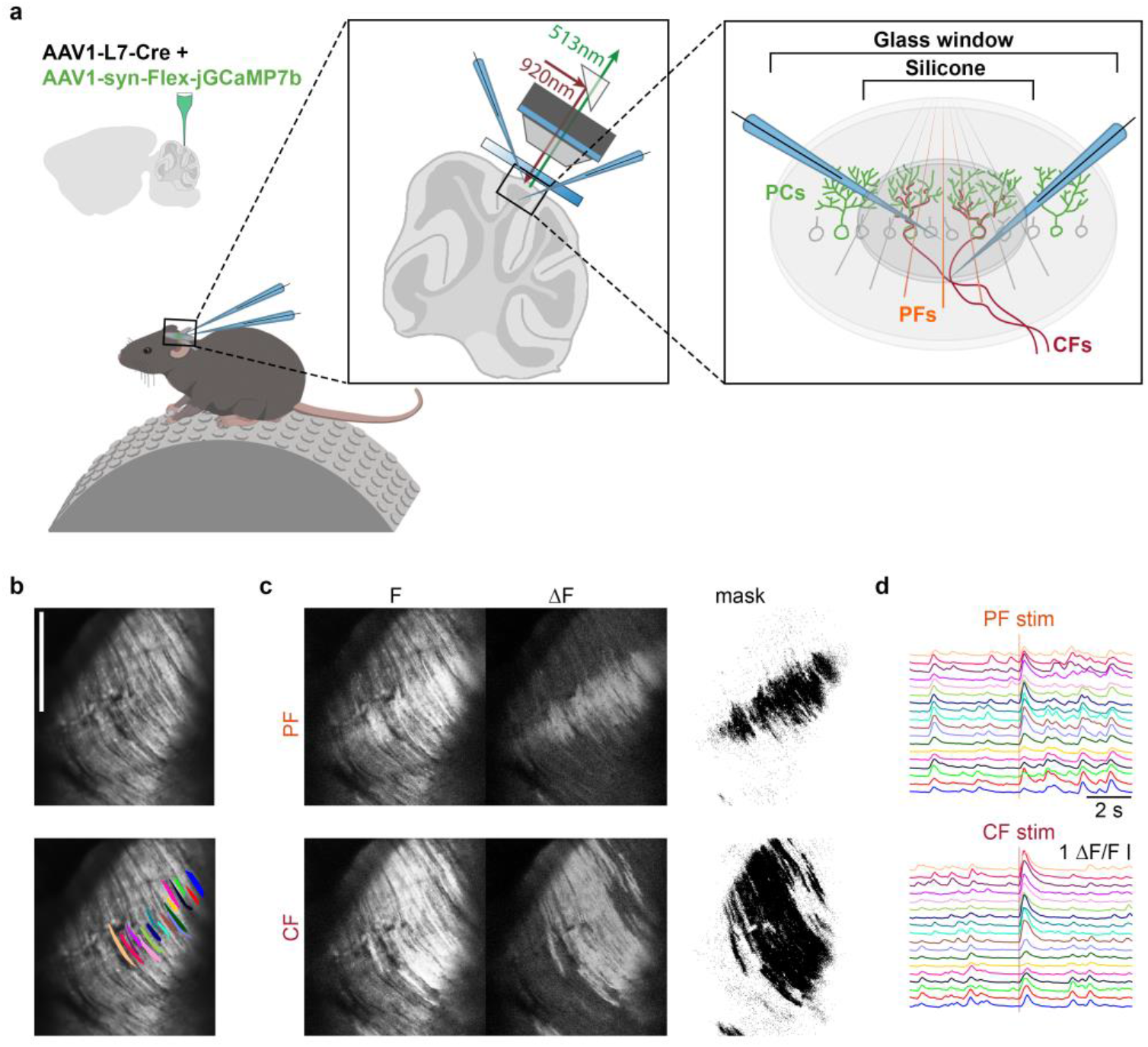
Electrical stimulation evokes PF or CF-mediated calcium transients in PC dendrites in awake mice. **a** Schematic of the experimental setup illustrating a calcium signal recording in cerebellar PCs. PC–specific jGCaMP7b expression was achieved via L7-Cre–dependent expression following AAV vector microinjection into the cerebellar cortex (Crus I). A customized glass window was implanted over Crus I, allowing two-photon imaging of calcium responses. Two microelectrodes were inserted through a silicone access port in the center of the window, where one was used to stimulate a bundle of PFs, and the other, positioned deeper, to stimulate the CF input. **b** The top panel shows a representative two-photon calcium image, and the bottom panel displays the same image overlaid with example regions of interest (ROIs). **c** Visualization of PF- and CF-responsive areas within the same field of view shown in (**b**). From left to right, average fluorescence (F), ΔF (F - F0, where F0 is defined as 20th percentile of the entire fluorescence trace), and a response mask generated using an arbitrary threshold within 300ms following PF (top) or CF (bottom) electrical stimulation (8pulses at 100Hz). **d** Representative calcium signals from the ROIs shown in (**b**). Only PCs that responded to both PF and CF stimulation are included.

Calcium signals control cerebellar synaptic plasticity; therefore, it is important to determine whether CF activity locally contributes to calcium signals at PF synapses. Spines on fine dendritic branches are exclusively contacted by PF synapses^18^. Nevertheless, CF stimulation evokes calcium transients in these spines as demonstrated *in vitro*^17^ as well as *in vivo*^19^. Under spontaneous (non-evoked) conditions, calcium signals can be measured in spines on fine dendritic branches (Fig. 2a-2b). These signals show a higher correlation when measured between spines located on the same dendrite than those on different dendrites (Fig. 2c), suggesting that spines located on the same dendrite form a functional signaling unit. This high within-dendrite coherence further supports the notion that these calcium signals reflect activity of one CF input.

**Fig. 2.**
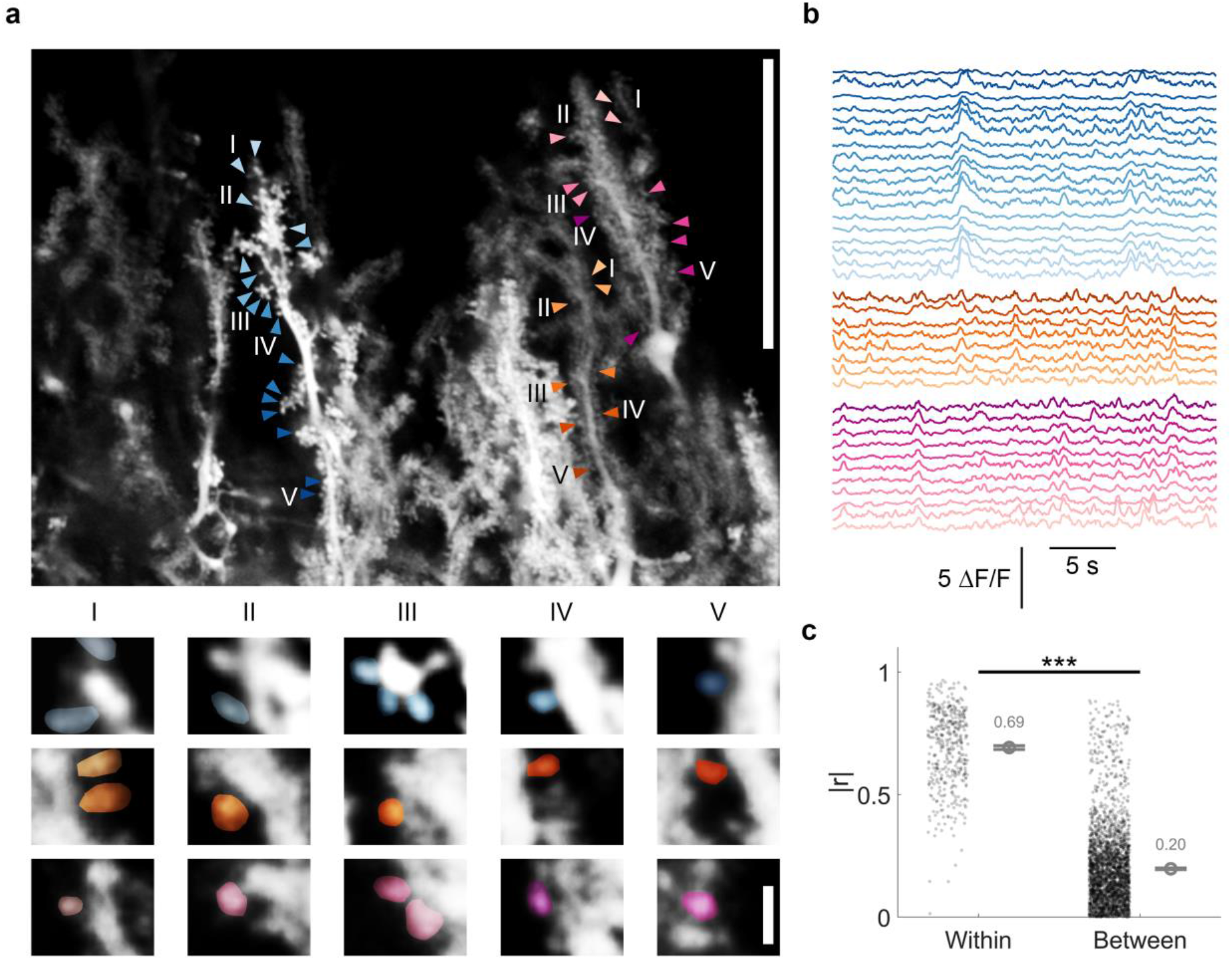
Two-photon calcium imaging shows PC spine signals. **a** Representative two-photon calcium image (top), and magnified views of individual dendritic spines overlaid with example ROIs (bottom). Arrows of the same color but different gradients indicate distinct spines on the same dendrite. Five magnified views (I-V; bottom) are shown for each of the three dendrites (blue, orange and magenta), corresponding to the labels in the top panel. Scale bars, 50 μm (top) and 2 μm (bottom). **b** Spontaneous (non-evoked) calcium activity in dendritic spines from three PC dendrites, with colors corresponding to those shown in (**a**). **c** Comparison of the absolute Pearson’s correlation coefficients (|r|) between synapse pairs within the same dendrites (n = 444) and between dendrites (n = 2826). Spontaneous calcium signals were obtained from 89 synapses across 8 dendrites within the same field of view. (two-sample t-test, ^***^p < 0.001)

To further determine whether CF activity induces calcium influx in PF spines, we measured dendritic spine calcium signals in response to either PF or CF stimulation (8 pulses at 100Hz each). The spines were selected from regions where PF and CF stimulus fields overlapped, and the majority exhibited robust calcium responses to PF stimulation (Fig. 3a, left), confirming that these spines are indeed contacted by PF synapses, as PF inputs typically generate highly localized calcium transients. Consistent with our observations in Fig. 2, CF stimulation also evoked calcium transients in the same PF spines (Fig. 3a, right). Because synaptic plasticity critically depends on calcium dynamics, we further characterized these responses to determine whether spine calcium signals differentially encode PF versus CF stimulation. First, CF responses exhibited significantly higher peak amplitudes (Fig. 3b), and lower trial-to-trial variability, as quantified by the coefficient of variation (CV; Fig. 3c), than PF responses, suggesting a stronger and more reliable CF-evoked response. Second, CF-evoked calcium transients show shorter rise-time kinetics (Fig. 3d) than those evoked by PF stimulation, consistent with previous observations of spontaneous dendritic calcium events^20^. Both PF and CF stimulation reliably evoked calcium responses, with response probabilities close to 1, although the probability of PF responses is slightly lower. The difference was not statistically significant (Fig. 3e). This result suggests that CF stimulation not only directly generates calcium influx in PF spines, but it may also evoke more reliable calcium responses at PF synapses even than PF stimulation itself. Because calcium responses are both detectable and distinct within individual spines, their differential dynamics are likely decoded by spine-located calcium sensors, particularly calcium/calmodulin-dependent kinase II (CaMKII)^21,22^, to regulate synaptic plasticity.

**Fig. 3.**
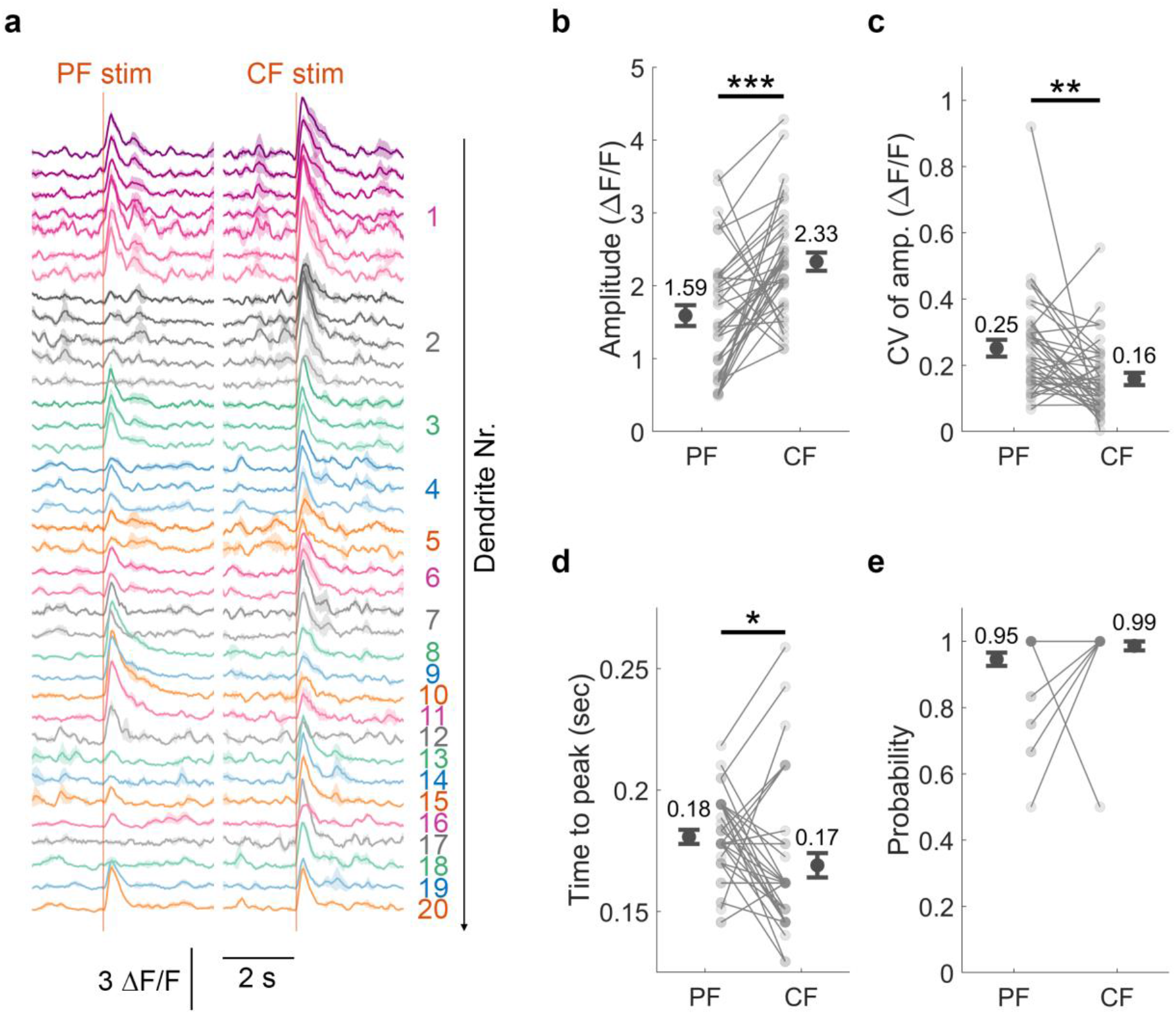
CF stimulation induces calcium transients at PC spines. **a** Calcium response (mean ± SEM) to PF or CF stimulation (8pulses at 100Hz for both). Signals were recorded from 37 spines across 20 dendrites in 5 mice. From top to bottom (Dendrite 1 to Dendrite 20), consecutive traces with the same color but different color depth represent individual spines belonging to the same dendrite. **b-e** Quantification of response parameters shown in (**a**): mean amplitude across trials (**b**), coefficient of variation (CV) of amplitude (**c**), time to peak from stimulation onset (**d**), and response probability (**e**). Small gray dots represent individual synapses, with lines connecting data from the same spine. Large black circles indicate the means ± SEMs. (paired t-test, ^*^p < 0.05; ^**^p < 0.01;^***^p < 0.001).

### Coincidence in PF and CF signaling does not drive LTD induction

To test whether coincident PF+CF activity evokes LTD in the intact mouse brain *in vivo* as it does *in vitro*, we used the same two-photon imaging strategy paired with electrical stimulation as described above. In these long-term recordings (>1h), however, we did not record from individual spines, but instead measured a response from the distal, fine dendritic branches spanning the full individual PC within the imaging plane. PF-evoked calcium transients collected this way vary in amplitude along with synaptic input strength and, although they are also influenced by intrinsic excitability, reflect synaptic plasticity well^23^.

The complete recording protocol consisted of baseline PF response measurements, followed by an induction protocol pairing PF and CF stimulation for ≥1min (see methods), and subsequent assessment of plasticity by measuring PF responses again (Fig. 4a). We first applied a simple pairing protocol (Fig. 4b), in which a single PF pulse was paired with a brief CF burst (4 pulses at 100Hz). Despite the efficacy of this protocol (and other similar ones) for LTD induction *in vitro*^7^, it did not cause LTD in our *in vivo* recordings (Fig. 4d-4e). Calcium transients evoked during the first 10 sec of the induction protocol show a regular series of responses that did not fade over time (Fig. 4b). Of note, application of the same PF stimulus pattern without CF stimulation causes long-term potentiation (LTP) of PF-evoked responses^23^, similar to earlier findings from cerebellar slices^7^. Even when a short PF burst (3 pulses at 50Hz) was applied instead of a single pulse and paired with the same CF burst (4 pulses at 100Hz), LTD was not induced (Fig. 4c, 4f, and 4g), despite the larger calcium responses observed during the induction protocol compared to single-pulse PF stimulation (Fig. 4c). Considering the heterogeneity in plasticity outcomes across individual PCs, we categorized each neuron into LTP, no-change, LTD outcome groups using the normalized baseline ± 15% change (1 ± 0.15) as criterion (right side of Fig. 4e and 4g). We observed that most cells (78% and 85%) fell into the no-change group under both induction protocols. These findings suggest that coincident PF+CF activation does not effectively drive LTD in the majority of PCs. Following the late recording period, a second induction protocol was applied using the same PF stimulation paired with four CF pulses at 333Hz, followed by two additional recording periods (Supplementary Fig. 1). A transient LTD was observed immediately after the induction protocol (early phase) with single-pulse PF stimulation, whereas no effect was observed after the short-burst protocol.

**Fig. 4.**
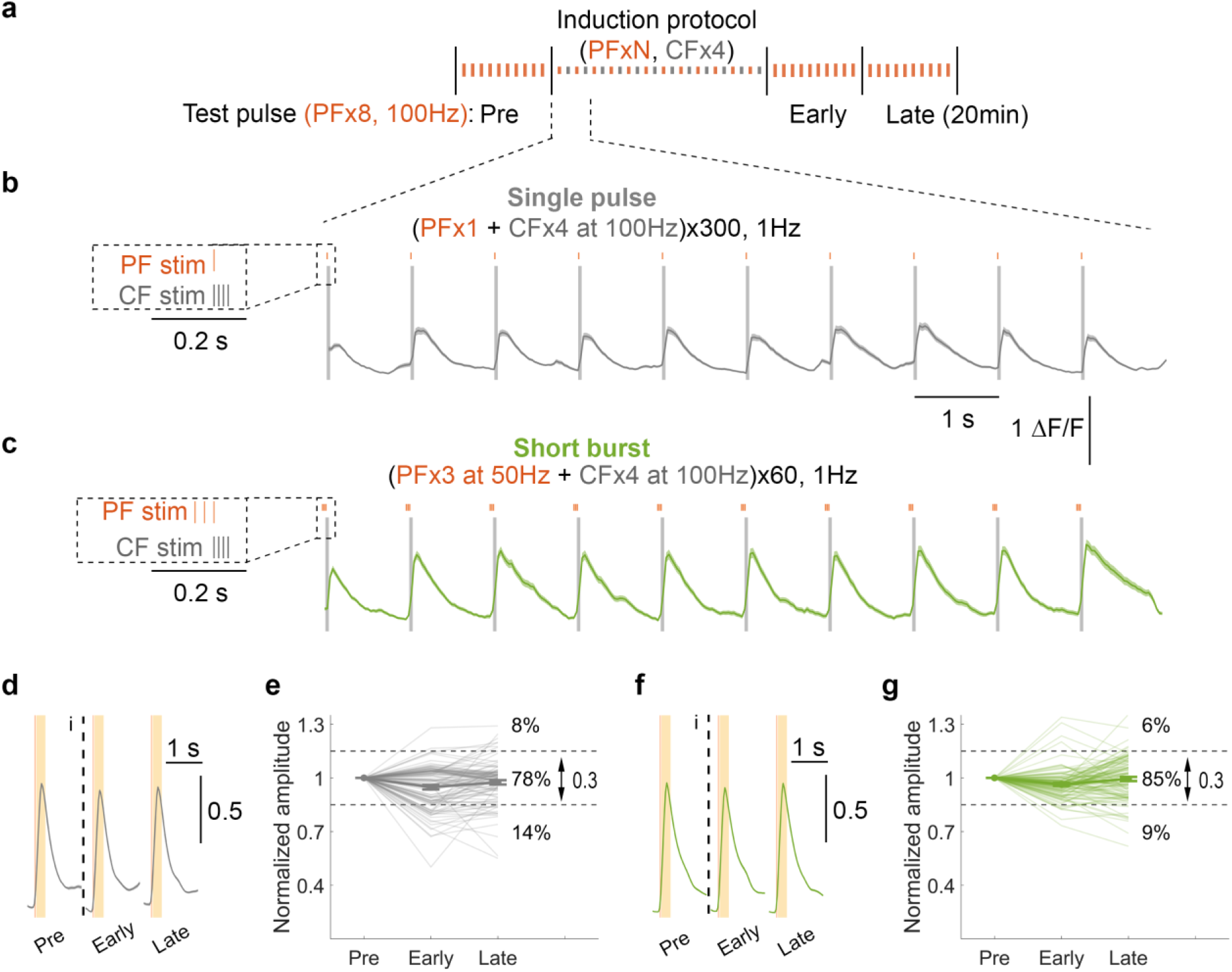
A single PF pulse or brief PF burst followed by a CF burst does not induce LTD. **a** Schematic illustration of experimental protocols. Baseline PF responses in PCs were recorded for 20 min, followed by the induction protocol and two subsequent 20-min recording periods (early and late). For the different induction protocols, PF stimulation parameters were varied as specified in panels **b** and **c**, and paired them with four-pulse CF burst (0.3ms per pulse) delivered at 100Hz. **b-c** Average calcium signals ± SEM during the first 10 sec of induction protocol, along with the magnifications of the stimulation protocols at left side. In (**b**), the protocol consists of a single PF pulse followed by CFx4 at 100Hz. In (**c**), the protocol consists of a short burst of PF stimulation (3xPF at 50Hz) followed by CFx4 at 100Hz. In both protocols, the last PF pulse was synchronized with the first CF pulse. The protocols were repeated 300 times at 1Hz for (**b**) and 60 times for (**c**). **d, f** Normalized average calcium signals ± SEM across different recording phases. Signals were normalized by the mean amplitude of pre-tetanus calcium transients (y-axis denotes normalized value units). Following the test pulse (PFx8 at 100Hz), the yellow shaded area (50-300ms) indicates the analysis window for the amplitude estimation. **e, g** Peak amplitude normalized to baseline (pre). Thin lines represent individual cells, and thick line indicate the mean ± SEM (values are provided in Supplementary Table 1). Neurons were classified into LTP, no-change, or LTD groups using a threshold, which is 15% of the normalized baseline amplitudes. Specifically, neurons were defined as LTP if the late-phase amplitude exceeded baseline + 0.15, no-change if within baseline ± 0.15, and LTD if below baseline – 0.15. Differences in amplitude across phases were assessed using one-way ANOVA (**e**, F(4, 505) = [8.2], p < 0.001, 102 cells/5 mice; **g**, F(4, 535) = [2.54], p < 0.039, 108 cells/4 mice; same statistical test applied to Supplementary Fig. 1e and 1g, respectively) followed by Tukey’s HSD post-hoc test (^*^p < 0.05; ^**^p < 0.01; ^***^p < 0.001).

### Preceding and ramping PF activity patterns promote LTD

To test whether preceding PF activity enables LTD, we first applied a tonic PF burst consisting of five pulses at 10Hz (Fig. 5b), which was followed by a CF burst (4 pulses at 100Hz). The calcium traces recorded during the first 10 sec of stimulation reveal that the five PF pulses preceding the CF burst caused small calcium transients that in some responses during the stimulus train summed up to reach higher amplitudes than the individual ones (Fig. 5b). In contrast to the previous two induction protocols shown on Fig. 4, this stimulation elicited LTD that was recorded after tetanization (Fig. 5d-5e).

**Fig. 5.**
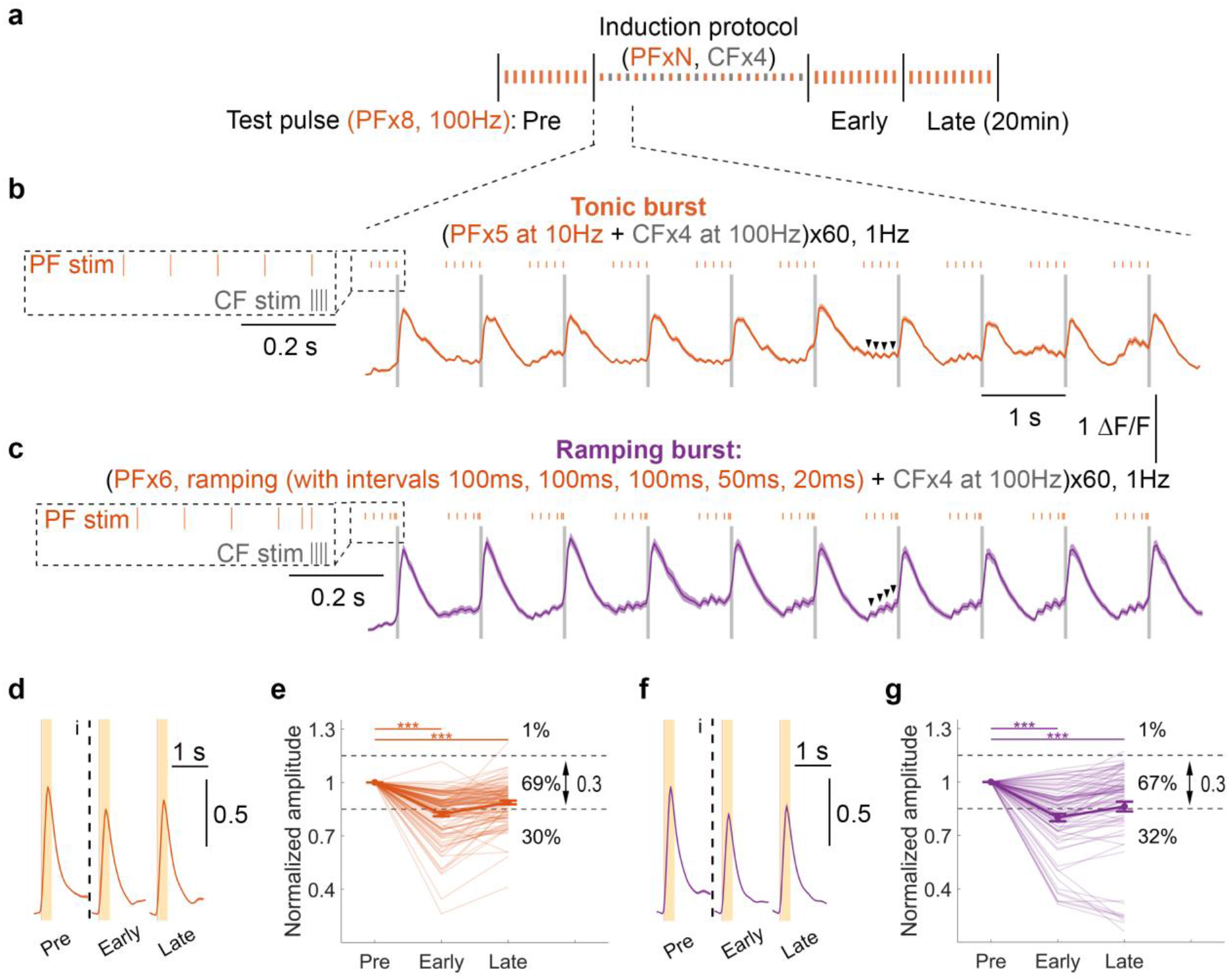
A tonic or ramping PF burst followed by a CF burst induces LTD. **a** Schematic illustration of experimental protocols. Baseline PF responses in PCs were recorded for 20 min, followed by the induction protocol and two subsequent 20-min recording periods (early and late). For the different induction protocols, PF stimulation parameters were varied as specified in panels **b** and **c**, and paired them with four-pulse CF burst (0.3ms per pulse) delivered at 100Hz. **b-c** Average calcium signals ± SEM during the first 10 sec of induction protocol, along with the magnifications of the stimulation protocols at left side. In (**b**), the protocol consists of a tonic burst of PF stimulation (5xPF at 10Hz) followed by CFx4 at 100Hz. In (**c**), the protocol consists of a ramping PF burst with intervals of 100, 100, 100, 50, 20ms followed by CFx4 at 100Hz. In both protocols, the last PF pulse was synchronized with the first CF pulse. In both cases, protocols were repeated 60 times at 1Hz. Arrows denote small calcium transients evoked by individual PF pulses. **d, f** Normalized average calcium signals ± SEM across different recording phases. Signals were normalized by the mean amplitude of pre-tetanus calcium transients (y-axis denotes normalized value units). Following the test pulse (PFx8 at 100Hz), the yellow shaded area (50-300ms) indicates the analysis window for the amplitude estimation. **e, g** Peak amplitude normalized to baseline (pre). Thin lines represent individual cells, and thick line indicate the mean ± SEM (values are provided in Supplementary Table 1). Neurons were classified into LTP, no-change, or LTD groups using a threshold, which is 15% of the normalized baseline amplitudes. Specifically, neurons were defined as LTP if the late-phase amplitude exceeded baseline + 0.15, no-change if within baseline ± 0.15, and LTD if below baseline – 0.15. Differences in amplitude across phases were assessed using one-way ANOVA (**e**, F(4, 570) = [34.76], p < 0.001, 115 cells/6 mice; **g**, F(4, 405) = [36.18], p < 0.001, 82 cells/5 mice; same statistical test applied to Supplementary Fig. 2e and 2g, respectively) followed by Tukey’s HSD post-hoc test (^*^p < 0.05; ^**^p < 0.01; ^***^p < 0.001).

To specifically test the efficacy of PF ramping, we applied a protocol, in which six PF pulses were applied before the same CF burst (Fig. 5c). This protocol consisted of three PF pulses at 10Hz, followed by one after 50ms (20Hz) and another after 20ms (50Hz), resulting in a 370ms ramping pattern. The calcium traces show robust calcium build-up during the ramping period that did not occur as reliably during the non-ramping tonic PF burst. This ramping protocol resulted in LTD (Fig. 5f-5g) that was indistinguishable from that observed with tonic preceding PF activity. However, when the respective stimulus patterns were applied a second time after 40min—this time with a higher frequency CF burst (4 pulses at 333Hz; Supplementary Fig. 2)—the ramping protocol further enhanced the magnitude of LTD (Supplementary Fig. 2f-2g), whereas the non-ramping protocol produced only a transient early-phase depression that barely reached statistical significance (Supplementary Fig. 2d-2e).

By quantifying neurons exhibiting different plasticity effects, we observed a clear shift toward LTD after both tonic and ramping induction protocols, reflected by a decrease in the LTP and no-change groups accompanied by an increase in neurons exhibiting LTD (Fig. 5e and 5g). Moreover, in neurons exhibiting LTD, tonic and ramping PF activity caused larger LTD amplitudes than those induced by coincident PF+CF stimulation (Fig. 4).

### Preceding PF-evoked calcium elevation determines LTD

By summarizing the changes in amplitudes and the proportion of PC populations exhibiting each plasticity outcome, we found that only preceding PF activity paired with a CF burst reliably induces LTD (Fig. 6a-6c). To understand how this plasticity outcome is determined by the calcium signaling during the induction protocol, we realigned and superimposed the calcium traces for each PF+CF stimulation pattern (Fig. 6d). From these traces, a set of parameters was quantified to characterize the calcium waveform (Fig. 6e), including Δbaseline (Fig. 6f), the area under curve (AUC; Fig. 6g), as well as relative and absolute amplitudes (Fig. 6h and 6i, respectively).

**Fig. 6.**
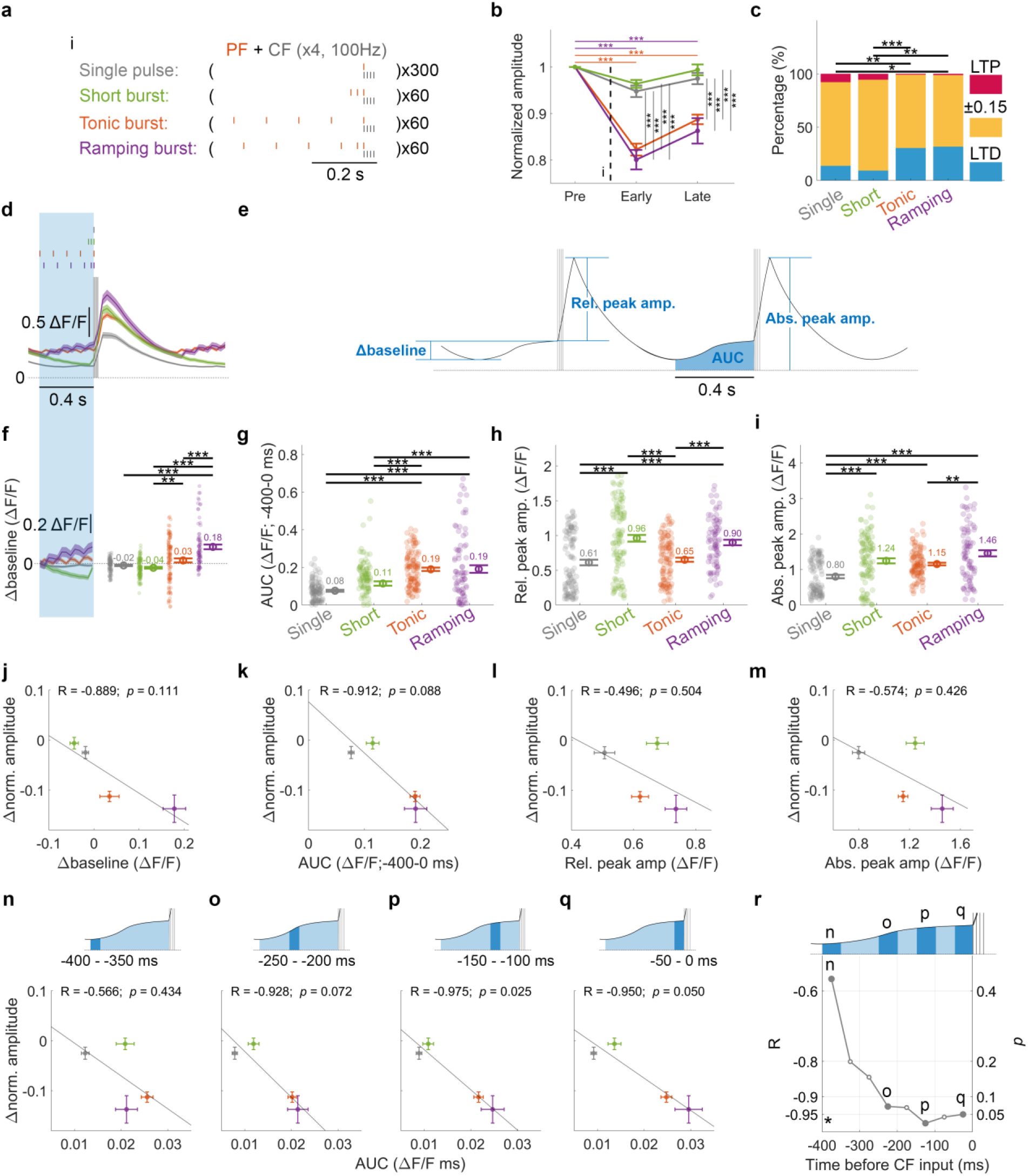
Prolonged calcium baseline elevation, not only the peak amplitude, facilitates LTD induction. **a** Schematic illustration of induction protocol in each experimental setup, aligned by the last PF pulse which coincides with the first CF pulse. **b** Peak amplitude before and after the induction protocol, normalized by the baseline value (pre). Statistical differences were assessed using Two-way ANOVA (F_protocol_[3,1209] = 51.95, p < 0.001; F_time_[2,1209] = 98.41, p < 0.001; F_protocol x time_[6,1209] = 13.73, p < 0.001; single pulse, 102 cells/5 mice; short burst, 108 cells/4 mice; tonic burst, 115 cells/6 mice; ramping burst, 82 cells/5 mice) followed by Tukey’s HSD post-hoc test (^*^p < 0.05; ^**^p < 0.01; ^***^p < 0.001). **c** Percentage distribution of LTP, no-change (labeled as “±0.15” to indicate the neurons fall in 1 ±0.15 range), and LTD neurons across four experimental conditions. Differences in the distributions were assessed using a chi-square test (X^2^[6, N = 407] = 31.026; p <0.001) followed by Bonferroni post-hoc comparisons (^***^p < 0.001). **d** Average calcium signals ± SEM during the first 10 seconds of the induction protocol. Bursts 2-9 were realigned based on the onset of CF stimulation. The blue shaded area, encompassing the temporal range of all different PF stimulation protocols, denotes the analysis window (-400 to 0 ms relative to CF burst onset) for calculating Δbaseline and the area under curve (AUC), as detailed in the next panel. Because repetitive PF tetanus progressively elevates the calcium baseline, F0 for this analysis was defined as the 20th percentile of the final sweep during the pre-phase recording. **e** Schematic illustrating how different response parameters were measured for the tetanus protocols. Δbaseline represents the fluorescence change within the PF analysis window (-400 to 0ms relative to the onset of CF stimulation). AUC represents the area under the curve within this window. Relative amplitude is defined as the peak amplitude relative to CF burst onset, whereas absolute amplitude is the peak amplitude relative to 0 ΔF/F0. **f-i** Individual values alongside with mean ± SEM of response parameters for the tetanus protocols. (**f**) Δbaseline (F[3, 403] = 30.56, p < 0.001), (**g**) AUC (F[3, 403] = 24.37, p < 0.001), (**h**) relative peak amplitude (F[3, 403] = 19.18, p < 0.001), and (**i**) absolute peak amplitude (F[3, 403] = 18.97, p < 0.001), where statistical differences were assessed using One-way ANOVA followed by post hoc comparisons using Tukey’s HSD. **j-m** Changes in PF response amplitudes during the late phase relative to the pre-phase baseline (**b**) are plotted against different response parameters of the tetanus protocols (**f-i**). The relationships were assessed using Pearson’s correlation coefficients (R), shown at the top of each plot along with corresponding p values. **n-q** Similar to **k**, changes in PF response amplitudes during the late phase relative to the pre-phase baseline (**b**) are plotted against AUC calculated within different time windows preceding CF stimulation. Relationships were assessed using Pearson’s correlation coefficients (R), shown at the top of each plot along with corresponding p values. **r** Pearson’s correlation coefficients (left axis) and the corresponding p values (right axis), derived from **n-q** and all other 50ms windows across 400ms period prior to CF stimulation, plotted as a function of time windows. The asterisk marks the line representing R = -0.95 and p = 0.05.

Δbaseline denotes the baseline change measured within a 400ms time window preceding the onset of the CF burst, covering the duration of the longest PF burst (i.e., tonic). This time window falls within the range commonly used in delayed-response behavioral assays that robustly engage cerebellar learning^24-27^. However, it does not fully capture the temporal scale of ramping activity observed under physiological conditions, where ramping activity preceding reward in conditioning experiments can extend for seconds^11,12,28^. Without preceding PF activity, calcium signals returned to baseline after the previous CF burst in the single-pulse and short-burst conditions (Fig. 6d and 6f). In contrast, in the other two protocols, calcium levels remained at an elevated level or progressively accumulated before the CF-evoked calcium transients. This elevated calcium baseline explains the higher AUC observed during tonic and ramping PF stimulation compared with the other conditions (Fig. 6g).

In contrast, both relative and absolute amplitudes (Fig. 6h-6i) during the CF co-stimulation are primarily enhanced by PF activity occurring close in time to the CF input, rather than by prolonged preceding PF activity. This is reflected in the short PF burst generating significantly higher amplitudes than single pulse stimulation in both amplitude metrics, and even higher amplitudes than tonic PF burst, although the latter difference did not reach statistical significance for the absolute amplitude. The relative amplitude (Fig. 6h) does not differ significantly between the tonic and single-pulse protocols, suggesting that the prolonged preceding low-frequency PF activity does not alter the peak calcium level resulting from the subsequent CF co-stimulation.

To further examine the relationship between calcium dynamics and the resulting synaptic plasticity, we plotted the relative change in PF response amplitude between the late and pre phases against each parameter (Fig. 6j-6m). Among the tested metrics, the AUC over 400ms period preceding CF stimulation best predicted LTD magnitude, compared with Δbaseline as well as both relative and absolute peak amplitudes. These results suggest that prolonged PF-evoked calcium elevation prior to CF input is a key determinant of LTD induction, rather than supralinear calcium signals induced by PF and CF coincidence.

To examine the temporal profile of the AUC-LTD relationship, we subdivided the 400ms preceding period into eight 50ms time bins, and correlated the AUC of each time bin with LTD magnitude. Four representative bins are shown (Fig. 6n-6q), along with a summary of Pearson’s correlation coefficients and the corresponding p values across all bins (Fig. 6r). The negative correlation gradually strengthened from -400ms toward CF onset, with the AUC in the -150 to - 100ms window best predicting LTD magnitude. From this time window onward, significant correlation emerged, aligning with calcium accumulation in tonic and ramping conditions.

## Discussion

Our study is the first to investigate how temporal relationships between PF and CF activity promote cerebellar LTD in an intact, awake animal preparation. The initial *in vivo* study describing LTD was conducted in rabbits by Masao Ito and colleagues, but the stimuli to the vestibular nerve and inferior olive, respectively, were applied continuously rather than in discrete patterns, preventing analysis of preferred temporal relationships^4^. More recent *in vivo* studies using delayed reward conditioning have revealed prolonged PF ramping activity between the initiation of expectation and reward-evoked CF input^11,12^, thereby introducing a temporal offset between these two pathways. However, how such temporal relationships shape synaptic plasticity—or, in other words, how synaptic plasticity encodes this temporal offset—remains unexplored. Our finding that this ramping PF activity, rather than coincident PF activity, drives LTD provides new insight into how synaptic plasticity encodes temporal differences in the cerebellum. Reminiscent of a ‘leaky integrator^29^, prolonged low-amplitude calcium signaling appears to facilitate LTD induction. Our spine recordings further revealed that PF and CF stimulation generate distinct calcium dynamics within PF synapses (Fig. 3), allowing calcium integrators such as CaMKII to differentially regulate synaptic plasticity. However, whether such mechanism involves a shift in the LTD threshold, as previously shown for low-versus high-frequency PF tetanization^22^, remains to be determined.

When interpreting our findings, it needs to be kept in mind that different cerebellar regions might show different plasticity rules that are optimized for specific tasks^8^. Our recordings were restricted to PCs located in Crus I, and it is possible that other lobules/areas would reveal different temporal pattern preferences. With the specific focus on Crus I, it is remarkable how distinct the plasticity outcomes are between protocols using preceding PF activity, and those using only coincident stimulation. Near-coincidence in PF and CF signaling has been seen, since the foundational theoretical work of David Marr (1969)^2^, as a prerequisite for cerebellar plasticity and motor adaptation. The LTD induction protocols designed to mimic this near-coincidence are Hebbian regarding the temporal precision and use of very short time intervals. In contrast, the protocols using preceding and ramping activity resemble behavioral timescale synaptic plasticity (BTSP)^30^ in that they integrate activity at synaptic inputs over the range of hundreds of milliseconds to seconds rather than a few milliseconds.

This timescale highlights a different, more recently appreciated aspect of cerebellar function: predictive coding. Ramping activity may signify the encoding of a sensory expectation, e.g. for a reward^11,12^. It is currently not known to what degree cerebellar circuits play a role in reward signaling *per se*, but reward constitutes a sensory event of relevance to the orientation of the animal in spacetime.

Another important aspect of using ramping activity is its role in tracking and learning of time intervals. During delayed reward tasks, PCs in Crus I and II exhibit ramping simple spike activity lasting up to 5 seconds^11^. In another example, granule cells in vermis (lobule VI) exhibit ramping activity over 2 seconds^12^. Even without reward components, ramping activity is broadly observed in teleost granule cells and can span windows of up to 21 seconds^31^. Together, these findings suggest that ramping inputs to PCs are widespread, with the observed time scales likely limited by experimental design. To examine how the duration of PF activity may be encoded at PF synapses, Garcia-Garcia and colleagues predicted LTD magnitudes using a conventional coincidence-based plasticity rule^12^, in which LTD magnitude depends on the level of PF activity within a relatively narrow 150ms window preceding CF inputs. With ramping input patterns, even this coincidence-based model predicted that LTD magnitude correlates with ramping duration extending up to ∼2 seconds before reward. Based on this framework, our data revealed that LTD magnitude is directly determined by the temporal extent of preceding PF activity, providing a refined model that may enhance the precision of interval encoding in synaptic strength.

Plasticity operating at longer intervals may perhaps be as informative for motor gain adaptation as a counterpart that is exclusively governed by millisecond precision. In so far, a deviation from Marr’s expectation that plasticity driving inputs need to occur ‘*at about the same time*’ is not too much of a paradigm shift. While this may to some extent be true, it is only the preceding, ramping activity that allows another computational motif to play an essential role: anticipation and evaluative synaptic gain change under control of the supervising CF signal. In coincidence-based plasticity, the presence or absence of the CF signal assumes the role of a simple evaluative signal indicating whether the same event occurs, without the motif of anticipation. Only the longer time interval allows for anticipation, in its most simple form in classical conditioning, such as delay eyeblink conditioning, which critically depends on the cerebellum^32^.

## Data availability

The raw 2-photon imaging data are too large to be uploaded to an online repository but are available upon request. The processed source data generated in this study have been deposited in the Zenodo database https://doi.org/10.5281/zenodo.19489362. Source data are provided with this paper.

### Code availability

The code is written in MATLAB R2021b and publicly available on GitHub (https://github.com/tingfenglin-ac/CerebellumRampingPFandLTD).

## Acknowledgements

We thank members of the Hansel laboratory for insightful discussions. This study was supported by funding from the National Institute of Neurological Disorders and Stroke (NIH-NINDS Grant R21NS140563).

## Author Contributions Statement

T.-F.L. and C.H. conceived and designed the experiments. T.-F.L. performed the surgeries, habituation, collected the data, wrote the analysis code. T.-F.L. and C.H. interpreted the data and wrote the manuscript.

## Materials and Methods

### Mice

All animal experiments were approved and conducted in accordance with the regulations and guidelines of the Institutional Animal Care and Use Committee of the University of Chicago. All mice were wildtype with C57BL/6J background (The Jackson Laboratory, Bar Harbor, ME, USA). P80-P120 adult mice were selected for the two-photon recording performed in this study, and the necessary surgeries were conducted 2–3 weeks prior to the recording. The plasticity experiments were done in mice number of 5 for single-pulse, 4 for short-burst, 6 for tonic-burst and 5 for ramping-burst experiments. Spontaneous PF dendritic spine signals were recorded from a single mouse. PF and CF responses in dendritic spines were derived from a subset of plasticity experiment recordings (pre-phase only), using signals from spines that could be visually identified. No sex differences were observed in the reported measures.

### Cranial window implantation and jGCaMP7b expression

To enable in vivo calcium imaging, mice underwent surgical implantation of a cranial window combined with jGCaMP7b viral delivery 2-3 weeks before imaging sessions. All procedures, including headplate installation, viral injection, and cranial window placement, were completed within 2 hours. Anesthesia was induced and maintained with inhaled isoflurane. After the anesthesia induction, analgesia was applied with subcutaneous meloxicam (1–2 mg/kg) and buprenorphine (0.1 mg/kg), along with sterile saline (1 mL) for hydration. A toe-pinch test was applied to confirm adequate depth of anesthesia. Throughout the procedure, body temperature was maintained at 35–37 °C using a DC Temperature Controller System (FHC, Bowdoin, ME, USA).

Except for the anesthesia protocol, all surgical procedures followed those described in our previous publication^23^. Briefly, fur over the surgical site was removed, and then skin was disinfected with alternating applications of betadine and 70% ethanol three times. After excising the skin and connective tissue, a custom-made titanium headframe (H. E. Parmer Company, Nashville, TN, USA) was affixed to the skull using dental cement (Stoelting Co., Wood Dale, IL, USA). A circular craniotomy (4 mm diameter) was then made using a dental drill. The opening was centered at 2.5 mm laterally from the midline and 2.5 mm caudally from lambda. Following removal of the dura, lobules simplex, Crus I, and anterior Crus II of the cerebellum were exposed. The brain surface was gently cleaned to remove any residual debris and blood clots before placement of the cranial window. A two-layer glass window was then installed and secured over the craniotomy using C&B Metabond dental cement (Patterson Dental Company, Saint Paul, MN, USA). The window assembly consisted of a 5 mm diameter coverslip (Warner Instruments, Holliston, MA, USA, # CS-5R) adhered to a 4 mm diameter coverslip (Tower Optical Corp, Boynton Beach, FL, # 4540-0495) using Norland Optical Adhesive (Norland Products Inc., Jamesburg, NJ) cured using UV light. To enable subsequent PF and CF electrical stimulation, a 1.5 mm diameter hole was drilled at the center of the two-layered glass window and filled with transparent Kik-Sil silicone adhesive (World Precision Instruments, Sarasota, FL, USA).

After placement of the cranial window, a glass pipette (tip diameter of ∼300 μm) was introduced through the silicone access port for viral delivery. A total of 1800 nl of AAV virus mix was injected into two sites (900 nl per site) in medial and lateral Crus I (1.8 mm and 3.2 mm lateral to the midline, respectively). At each site, 450 nl was delivered at depths of 500 μm and 250 μm below the pial surface. To achieve sparse jGCaMP7b expression, the viral mixture contained 0.5% of full titer L7-Cre (AAV1.sL7.Cre.HA.WPRE.hGH.pA, Princeton Neuroscience Institute (PNI) Viral Core Facility, kindly provided by the lab of Dr. Samuel Wang at Princeton University), and 20% Cre-dependent jGCaMP7b (AAV-syn-FLEX-jGCaMP7b-WPRE; Addgene, #104493). After injection, the glass pipette was left in place for 5 min before being withdrawn.

### Animal habituation

Following viral injection, mice were monitored daily for 5-7 days to ensure post-operative recovery. Habituation training started once the animals resumed normal exploratory behavior and showed no signs of distress (e.g., trembling or vocalization). Habituation began with gentle handling. Once signs of handling-related anxiety diminished, typically within two days, the duration was increased from 15 min to 30 min. Handling was performed near the imaging setup so that mice could become familiar with the environment and apparatus. After two days of handling, mice were gradually introduced to a free-running treadmill. The duration of treadmill exposure was progressively extended according to their comfort levels, ranging from 10 minutes to 1.5 hours per day. Habituation continued for 1–2 weeks, until mice remained calmly on the treadmill for more than 1.5 hours without displaying aversive behaviors.

### Two-photon microscopy

Calcium imaging of jGCaMP7b was performed in Crus I of the right cerebellar hemisphere in head-fixed mice using a laser scanning two-photon microscope (Neurolabware, Los Angeles, CA, USA). Motor behavior was continuously monitored with a DALSA M640 CCD camera (Teledyne Technologies, Thousand Oaks, CA, USA), and imaging was initiated only when the mouse was stationary. Images were typically acquired at frame rate of 31 Hz with a frame size of 635 × 512 pixels using an 8KHz resonant scanning mirror. For recording of spontaneous spine activity, imaging was performed at a reduced frame rate of 15 Hz.

The calcium indicator jGCaMP7b was excited using a 920 nm laser source generated by a Mai Tai® DeepSee system (Spectra-Physics, Milpitas, CA, USA). Fluorescence signals were collected through a 16x water dipping objective (Nikon LWD 0.8NA, 3 mm WD) and detected using a GaAsP PMT (Hamamatsu Photonics, Shizuoka, Japan). Image acquisition and microscope control were managed using Scanbox software (Scanbox, Los Angeles, CA), which implemented 3-4× digital zoom for plasticity experiments and an 8x digital zoom for spine recording. To reduce ambient light contamination, a custom-built light shield was positioned around the objective and cranial window during imaging.

### Electrical PF and CF stimulation

To evoke PF or CF stimulation *in vivo*, two glass pipettes filled with ACSF were inserted through the silicone access port and positioned in the molecular layer and the granule cell layer or white matter, respectively. The first one was used to stimulate a bundle of PF fibers, while the second pipette delivered CF stimulation. PF stimulation was delivered more than 100 μm away along the mediolateral axis. The CF stimulation site was positioned below the Purkinje cell layer but was often shifted anteriorly or posteriorly (along the sagittal axis) relative to the imaged PCs. To locate the appropriate stimulation site, glass pipettes were controlled by a PatchStar motorized micromanipulator (Scientifica, Uckfield, UK) while identifying the resulting calcium responses using epifluorescence imaging. The PG4000A digital stimulator (Cygnus Technology, Delaware Water Gap, PA, USA) was used to generate the stimulation protocol, which triggered the SIU91A (Cygnus Technology) to deliver electrical pulses via TTL signals. The initiation of stimulation protocol was synchronized with the two-photon imaging using TTL signals sent from scanbox board.

Each experiment consisted of five recording phases and two induction protocols: a pre-tetanus baseline period, followed by the first induction protocol consisting of different PF stimulus paired with four CF burst at 100Hz, and then early and late post-tetanus recording periods (Fig. 4a and Fig. 5a). This sequence was followed by a second induction protocol using the same PF stimulation patterns paired with four CF burst at 333Hz, again followed by early and late post-tetanus periods (Supplementary Fig. 1a and Supplementary Fig. 2a). Only the first three recording phases are shown in the main figure (Fig. 4a and Fig. 5a). Each recording phase lasted 20 min and consists of 10–12 trials spaced approximately 1–2 min apart. During the recording phase, each trial lasted 10 seconds, and a test stimulus (eight pulses, 0.3 ms duration at 100 Hz) was delivered to a bundle of PFs at the 5 second mark. During the induction protocols, PF stimulation patterns included a single pulse (single-pulse), three pulses at 50Hz (short-burst), five pulses at 10Hz (tonic-burst), or six pulses with progressively shorter intervals (100, 100, 100, 50, 20ms; ramping-burst). Each PF pattern was followed by four pulses of CF delivered at either 100Hz or 333Hz, with the last PF pulse synchronized at the first CF pulse. The PF+CF pairing was repeated 60 times at 1Hz in most protocols (short-burst, tonic-burst and ramping burst), but 300 times in the single-pulse protocol to test whether increasing the number of pairings could drive LTD.

### Image analysis

The imaging analysis pipeline was described in our previous publication^23^. Unless otherwise specified, the analyses were performed using custom-made MATLAB R2017b scripts (MathWorks, Natick, MA, USA; see Code Availability). Specifically, motion correction was performed using a whole-frame cross-correlation script (derived from a script provided by Mark Sheffield, University of Chicago^33^). Regions of interest (ROIs) for different analyses were manually drawn using Fiji ImageJ. The ROIs were used to extract calcium fluorescence intensity, and then ΔF/F was calculated with F0 baseline defined as the 20th percentile of each fluorescence trace^25^. To analyze calcium signals recorded during the induction protocol, because the high-frequency induction protocol does not provide an appropriate baseline using its own 20th percentile value, F0 was instead defined as the 20th percentile of the fluorescence trace from the previous PF test trial during pre-tetanus recording phase. All ΔF/F traces were smoothed using a low-pass filter through a 5-frame moving window.

### Visualization and statistical analysis

To evaluate the plasticity effect, the peak amplitude of the PF response during each test trial was measured as the maximum value within a time window of 50–300ms after the onset of PF stimulation. To demonstrate the relative change of peak amplitudes, the average value from each recording phase of each PC was normalized by its own pre-phase baseline. To estimate the plasticity effects across recording phases, one-way ANOVA was performed (Fig. 4, Supplementary Fig. 1, Fig. 5, Supplementary Fig. 2), followed by post-hoc pairwise comparisons using Tukey’s HSD test. When the induction protocol was included as an additional factor, two-way ANOVA was performed (Fig. 6), followed by the same post-hoc test (Tukey’s HSD). The plasticity outcome of each PC after the first induction protocols (different PF stimulation patterns paired with 100Hz CF burst) was categorized as LTP, no-change, or LTD based on the normalized late post-tetanus value using a 1 ± 0.15 criterion: values >1.15 were classified as LTP, values between 0.85 and 1.15 as no-change, and values <0.85 as LTD (Fig. 4e, 4g, Fig. 5e, 5g, and Fig. 6c). The distributions of PCs across categories after different induction protocols were compared using chi-square test followed by Bonferroni post-hoc comparisons.

Calcium traces during the induction protocols (first 10 seconds only) are illustrated as average calcium signals ± SEM (Fig. 4b, c and Fig. 5b, c). The PF+CF stimulation patterns were repeated 10 times within this 10-second window, and the calcium responses from each repetition were realigned and averaged, excluding the first response, to better demonstrate the calcium dynamics evoked by the induction protocol (Fig. 6d). Differences in each calcium dynamic measurement during the induction protocols (Fig. 6f-6i) were assessed using One-way ANOVA followed by Tukey’s HSD post-hoc test. The relationship between these measurements (Fig. 6f-6i) and the resulting plasticity effects (late post-tetanus value − pre-tetanus value; Fig. 6b) was evaluated using Pearson’s correlation coefficient.

## Supplementary Information

**Supplementary Fig. 1.**
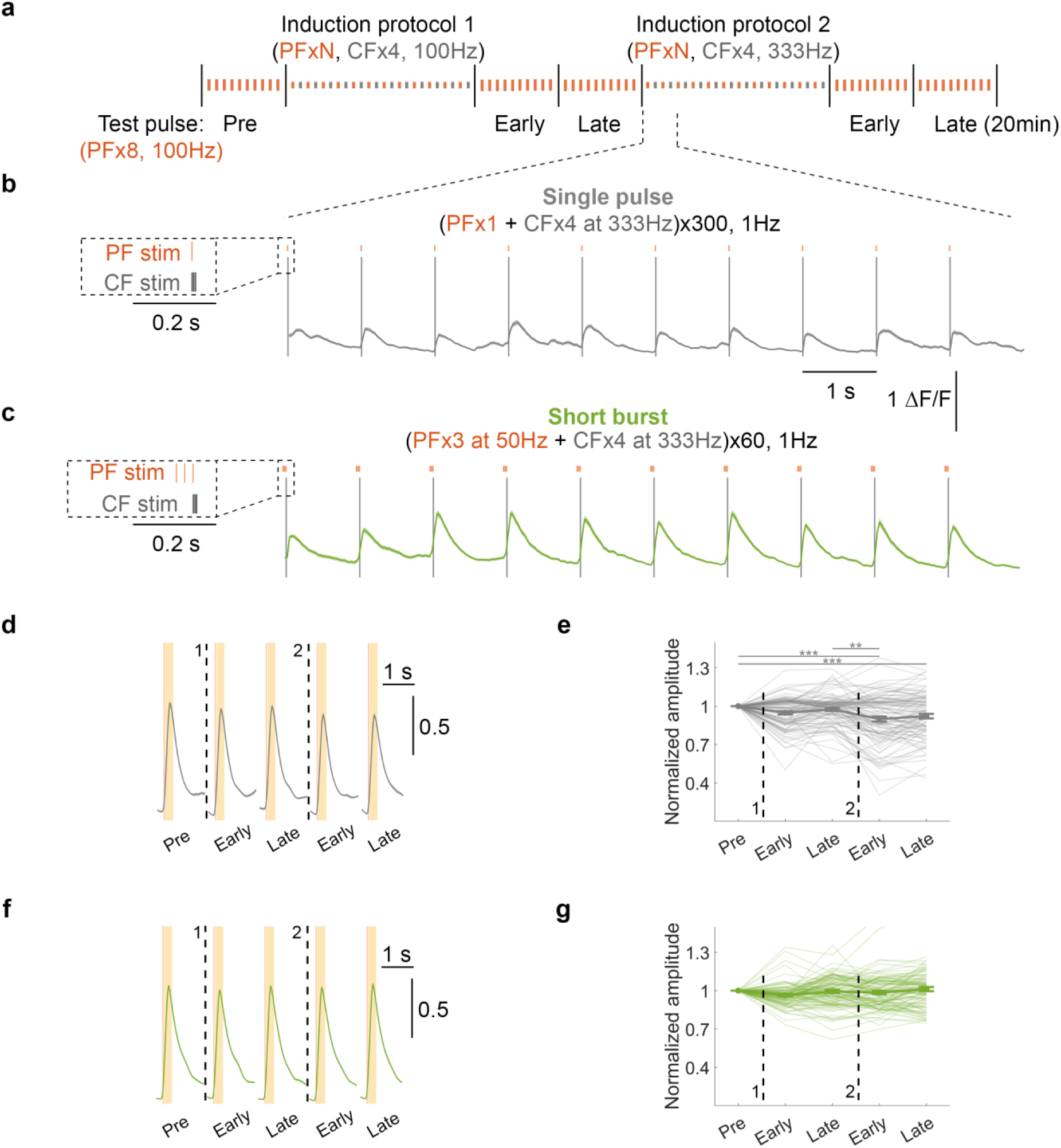
A further induction protocol with 333Hz CF burst. **a** Schematic illustration of experimental protocols. Baseline PF responses in PCs were recorded for 20 min, followed by the first induction protocol (1) and two 20-min recording periods (the same recording shown in **Fig. 4**). A second induction protocol (2) was then applied, followed by two recording periods. These two induction protocols consisted of PF stimulation (specified below), each paired with four CF pulses (0.3ms duration) delivered at 100Hz (1) or 333Hz (2). **b, c** Average calcium signals ± SEM during the first 10 sec of induction protocol (2), along with the magnifications of the corresponding stimulation protocols shown on the left. In (**b**), the protocol consists of a single PF pulse followed by CFx4 at 333Hz. In (**c**), the protocol consists of a short burst of PF stimulation (3xPF at 50Hz) followed by CFx4 at 333Hz. In these protocols, either the only PF pulse or the last PF pulse was synchronized with the first CF pulse. The protocols were repeated 300 times for (**b**) and 60 times for (**c**) at 1Hz. **d, f** Normalized average calcium signals ± SEM across different recording phases (the first three calcium transients are the same as Fig. 4d and 4f). Signals were normalized by the mean amplitude of pre-tetanus calcium transients (y-axis denotes normalized value units). Following the test pulse (PFx8 at 100Hz), the yellow shaded area (50-300ms) indicates the analysis window for the amplitude estimation. **e, g** Peak amplitude normalized by the baseline value (pre). Thin lines represent individual cells, and thick line indicate the mean ± SEM (values are provided in Supplementary Table 1). Statistical differences were assessed using One-way ANOVA (**e**, F(4, 505) = [8.2], p < 0.001, 102 cells/5 mice; **g**, F(4, 535) = [2.54], p < 0.039, 108 cells/4 mice; same statistical test applied to Fig. 4e and Fig. 4g, respectively) followed by Tukey’s HSD post-hoc test (^*^p < 0.05; ^**^p < 0.01; ^***^p < 0.001).

**Supplementary Fig. 2.**
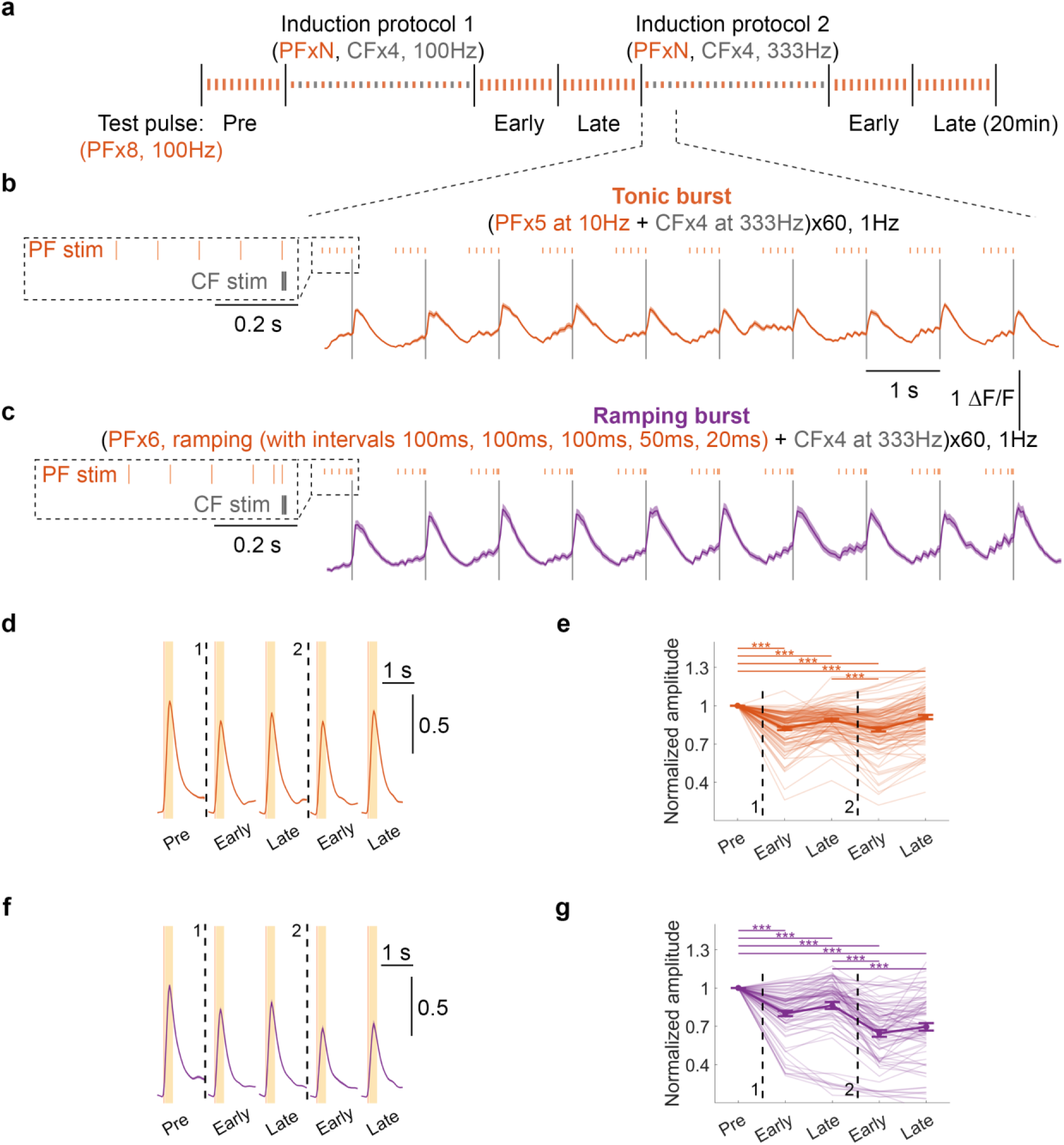
A ramping protocol with 333Hz CF burst leads to further LTD. **a** Schematic illustration of experimental protocols. Baseline PF responses in PCs were recorded for 20 min, followed by the first induction protocol (1) and two 20-min recording periods (the same recording shown in Fig. 5). A second induction protocol (2) was then applied, followed by two recording periods. These two induction protocols consisted of PF stimulation (specified below), each paired with four CF pulses (0.3ms duration) delivered at 100Hz (1) or 333Hz (2). **b-c** Average calcium signals ± SEM during the first 10 sec of induction protocol (2), along with the magnifications of the corresponding stimulation protocols shown on the left. In (**b**), the protocol consists of a tonic burst of PF stimulation (5xPF at 10Hz) followed by CFx4 at 333Hz. In (**c**), the protocol consists of a ramping PF burst with intervals of 100, 100, 100, 50, 20ms followed by CFx4 at 333Hz. In these protocols, the last PF pulse was synchronized with the first CF pulse. These protocols were repeated 60 times at 1Hz. **d, f** Normalized average calcium signals ± SEM across different recording phases (the first three calcium transients are the same as Fig. 5d and 5f). Signals were normalized by the mean amplitude of pre-tetanus calcium transients (y-axis denotes normalized value units). Following the test pulse (PFx8 at 100Hz), the yellow shaded area (50-300ms) indicates the analysis window for the amplitude estimation. **e, g** Peak amplitude normalized by the baseline value (pre). Thin lines represent individual cells, and thick line indicate the mean ± SEM (values are provided in Supplementary Table 1). Statistical differences were assessed using One-way ANOVA (**e**, F(4, 570) = [34.76], p < 0.001, 115 cells/6 mice; **g**, F(4, 405) = [36.18], p < 0.001, 82 cells/5 mice; same statistical test applied to Fig. 5e and Fig. 5g, respectively) followed by Tukey’s HSD post-hoc test (^*^p < 0.05; ^**^p < 0.01; ^***^p < 0.001).

**Supplementary Table 1.** Peak amplitude of the PF response across recording phases. Vaues are reported as mean ± SEM, corresponding to data shown in Fig. 4e and 4g; Supplementary Fig. 1e and 1g; Fig. 5e and 5g; and Supplementary Fig. 2e and 2g. From left to right, columns indicate baseline (Pre), the early (1) and late (1) phases following the first induction protocol, and the early (2) and late (2) phases following the second induction protocol. Each induction protocol consisted of PF stimulation (specified in the left column) paired with four CF pulses at either 100Hz (1) or 333Hz (2).

|  | **Pre** | **Early (1)** | **Late (1)** | **Early (2)** | **Late (2)** |
| --- | --- | --- | --- | --- | --- |
| **Single pulse** | 1.000 ± 0.000 | 0.948 ± 0.012 | 0.975 ± 0.012 | 0.900 ± 0.019 | 0.922 ± 0.018 |
| **Short burst** | 1.000 ± 0.000 | 0.964 ± 0.008 | 0.994 ± 0.012 | 0.986 ± 0.013 | 1.012 ± 0.016 |
| **Tonic** | 1.000 ± 0.000 | 0.822 ± 0.013 | 0.887 ± 0.010 | 0.815 ± 0.016 | 0.911 ± 0.017 |
| **Ramping** | 1.000 ± 0.000 | 0.801 ± 0.021 | 0.863 ± 0.027 | 0.644 ± 0.026 | 0.696 ± 0.029 |

